# ULK1-dependent phosphorylation of PKM2 antagonizes O-GlcNAcylation and inhibits the Warburg effect in breast cancer

**DOI:** 10.1101/2023.09.14.557850

**Authors:** Zibin Zhou, Xiyuan Zheng, Jianxin Zhao, Aiyun Yuan, Guangcan Shao, Bin Peng, Meng-Qiu Dong, Quan Xu, Xingzhi Xu, Jing Li

**Affiliations:** Beijing Key Laboratory of DNA Damage Response and College of Life Sciences, Capital Normal University, Beijing 100048, China; Guangdong Key Laboratory for Genome Stability & Disease Prevention and Carson International Cancer Center and Marshall Laboratory of Biomedical Engineering, Shenzhen University School of Medicine, Shenzhen, Guangdong 518060, China; National Institute of Biological Sciences, Tsinghua Institute of Multidisciplinary Biomedical Research, Tsinghua University, Beijing 102206, China; Department of Rehabilitation Medicine, Beijing Tsinghua Changgung Hospital, School of Clinical Medicine, Tsinghua University, Beijing 102218, China

**Keywords:** O-GlcNAcylation, ULK1, PKM2, breast cancer, Warburg effect

## Abstract

Pyruvate kinase M2 (PKM2) is a central metabolic enzyme driving the Warburg effect in tumor growth. Previous investigations have demonstrated that PKM2 is subject to O-linked β-N-acetylglucosamine (O-GlcNAc) modification, which is a nutrient-sensitive post-translational modification. Here we found that unc-51 like autophagy activating kinase 1 (ULK1), a glucose-sensitive kinase, interacts with PKM2 and phosphorylates PKM2 at Ser333. Ser333 phosphorylation antagonizes PKM2 O-GlcNAcylation, promotes its tetramer formation and enzymatic activity, and decreases its nuclear localization. By downregulating glucose consumption and lactate production, PKM2 pS333 attenuates the Warburg effect. Through mouse xenograft assays, we demonstrate that the phospho-deficient PKM2-S333A mutant promotes tumor growth *in vivo.* In conclusion, we identified a ULK1-PKM2-c-Myc axis in inhibiting breast cancer, and a glucose-sensitive phosphorylation of PKM2 in modulating the Warburg effect.

## Introduction

Facing rapid proliferation, cancer cells undergo metabolic reprogramming and shunt the consumed glucose to aerobic glycolysis, which is known as the Warburg effect (1). It enables cancer cells to proliferate in an uncontrolled fashion by providing increased ATP and biomass, and shifts the cellular metabolism from catabolic oxidative phosphorylation to anabolic aerobic glycolysis (2). Sitting at the heart of the Warburg effect is the pyruvate kinase M2 (PKM2) enzyme, which is a splicing variant of *PKM* (3). PKM1 and PKM2 differ only in one exon, but they are in sharp contrast to each other. PKM1 is expressed in most adult tissues, and exists only in tetramers. On the contrary, PKM2 is expressed during the embryonic stage, as well as in proliferative and cancer cells (2). PKM2 toggles between active tetramers and inactive dimers or monomers, so that its activity is finetuned (4). Functioning as the final rate-limiting enzyme, PKM converts phosphoenolpyruvate to pyruvate, thus its activity closely correlates with the carbon influx.

Due to the pivotal role of PKM2 in Warburg effect, it is both regulated by allosteric effectors and post-translational modifications (PTMs). Allosterically, PKM2 is activated by fructose 1,6-bisphosphate (FBP) and Serine availability (5,6). Post-translationally, modifications on PKM2 modulate its FBP-binding affinity, localization, stability and enzymatic activity: Lys-433 acetylation and Tyr-105 phosphorylation hamper the affinity between PKM2 and FBP (7,8); Ser-37 phosphorylation regulates PKM2 nuclear localization, where PKM2 exerts its nonmetabolic function in the Warburg effect (9); Lys-305 acetylation degrades PKM2 by the chaperone-mediated autophagy pathway (10), and PKM2 is also degraded by the ubiquitin E3 ligase OUT domain-containing ubiquitin aldehyde-binding protein 2 (OTUB2) (11); Met-239 oxidation sustains PKM2 tetramer formation for activation (12).

Of particular interest to us is the O-linked β-N-acetylglucosamine (O-GlcNAc) modification. Being the only intracellular mono-saccharide modification, O-GlcNAcylation occurs on protein Ser or Thr residues and functions as a nutrient rheostat to mediate cellular response to various stimuli (13). O-GlcNAcylation often crosstalks with phosphorylation, forming a yin-yang relationship in regulating both physiology and pathology (14). In the case of PKM2, it is first found to be O-GlcNAcylated at Thr-405 and Ser-406, which destabilizes PKM2 tetramers, leads to PKM2 nuclear translocation by promoting Ser-37 phosphorylation, and increases glucose consumption and promotes the Warburg effect (15). Later investigations further revealed that high glucose induces PKM2 O-GlcNAcylation at Ser-362 and Thr-365, which inhibits its PK activity and tetramer formation (16).

In our previous study of unc-51 like autophagy activating kinase 1 (ULK1) O-GlcNAcylation (17), we found that PKM2 is present in the ULK1 immunoprecipitates, raising the intriguing possibility that ULK1 could regulate PKM2. Herein we demonstrate that ULK1 phosphorylates PKM2 at Ser-333, which is downregulated when glucose is in surplus. Ser-333 phosphorylation antagonizes PKM2 O-GlcNAcylation, decreases PKM2 Ser-37 phosphorylation and its nuclear import, downregulates glucose consumption and lactate production, enhances PK activity by stabilizing tetramer formation. Further *in vivo* experiments showed that PKM2 Ser-333 phosphorylation attenuated breast cancer growth. Taken together, we identified a glucose-sensitive phosphorylation on PKM2 that antagonizes O-GlcNAcylation. Our findings suggest that the crosstalk between phosphorylation and O-GlcNAcylation may coordinate glucose status with cancer metabolism.

## Results

### ULK1 interacts with PKM2

In a recent investigation on PKM2 O-GlcNAcylation, PKM2 T405/S406 O-GlcNAcylation correlates with S37 phosphorylation, but PKM2 S37 mutation has no effect on O-GlcNAcylation (15). We reasoned that there could be other glucose-sensitive phosphorylation events that crosstalk with O-GlcNAcylation. To test this possibility, we used a pan phosphorylation antibody (pS/T) and examined PKM2 phosphorylation under different glucose concentrations (Fig. 1A). Remarkably, high glucose significantly downregulated the pS/T signal. In our previous investigation of ULK1 O-GlcNAcylation (17), PKM2 was present in the mass spectrometry (MS) analysis of ULK1 immunoprecipitates (Fig. 1B), suggesting that they could interact. As ULK1 is sensitive to glucose levels, we set out to investigate the potential interaction between ULK1 and PKM2. Cells were transfected with HA-PKM2 and Flag-ULK1 plasmids, and the lysates were immunoprecipitated (IPed) with anti-HA or anti-Flag antibodies. The results revealed that exogenous ULK1 and PKM2 co-IP reciprocally (Fig. 1C-D). Endogenous coIP was also observed between ULK1 and PKM2 (Fig. 1E). These results suggest that ULK1 binds with PKM2. We also examined the ULK1-PKM2 interaction when glucose is high (Fig. 1F-G). Remarkably, high glucose attenuated ULK1-PKM2 binding, correlating with decreased PKM2 phosphorylation under this condition (Fig. 1A). Taken together, we found that ULK1 interacts with PKM2 in a glucose-sensitive manner.

**Fig 1.**
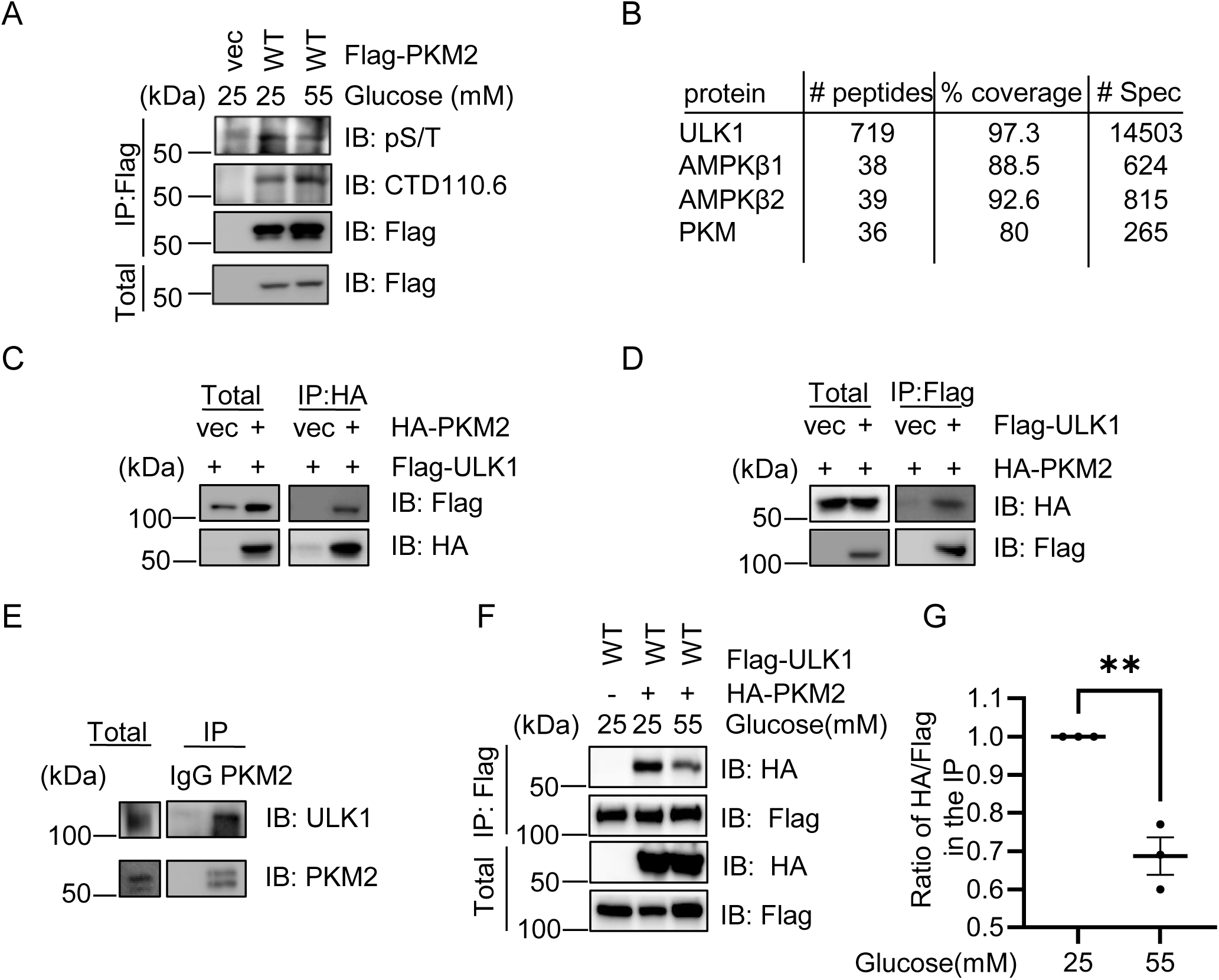
ULK1 interacts with PKM2. *A,* 293T cells were treated with glucose or mock treated. The cell extracts were then immunoprecipitated with anti-Flag antibodies and immunoblotted with the antibodies indicated. *B,* PKM is present in the ULK1 immunoprecipitates, as revealed by previous mass spectrometry analysis (17). *C* and *D*, co-immunoprecipitation between exogenous ULK1 and PKM2. 293T cells were transfected with FLAG-ULK1 and HA-PKM2 plasmids, and the cell lysates were subject to immunoprecipitation and immunoblotting experiments using the antibodies indicated. *E*, co-immunoprecipitation between endogenous ULK1 and PKM2. PKM2 immunoprecipitants were immunoblotted with anti-PKM2 and anti-ULK1 antibodies. *F,* High glucose decreases binding between PKM2 and ULK1. *G*, Quantitation of F. The statistical analysis in G was performed using Student’s *t*-test, **P < 0.01. All Western Blots were performed for at least three times.

### ULK1 phosphorylates PKM2 at S333

To explore the exact phosphorylation site on PKM2, *in vitro kinase* assays were carried out as previously described (18). The recombinant PKM2 proteins were efficiently phosphorylated by ULK1 (Fig. 2A). Then *in vitro kinase* products were analyzed by liquid chromatography-tandem MS (LC-MS/MS), and only Ser-333 conforms to the ULK1 consensus site (19) (Fig. 2B-C). We then generated recombinant GST-PKM2-S333A and used it in the *in vitro kinase* assay (Fig. 2A), and found that the phosphorylation signal was greatly attenuated.

**Fig 2.**
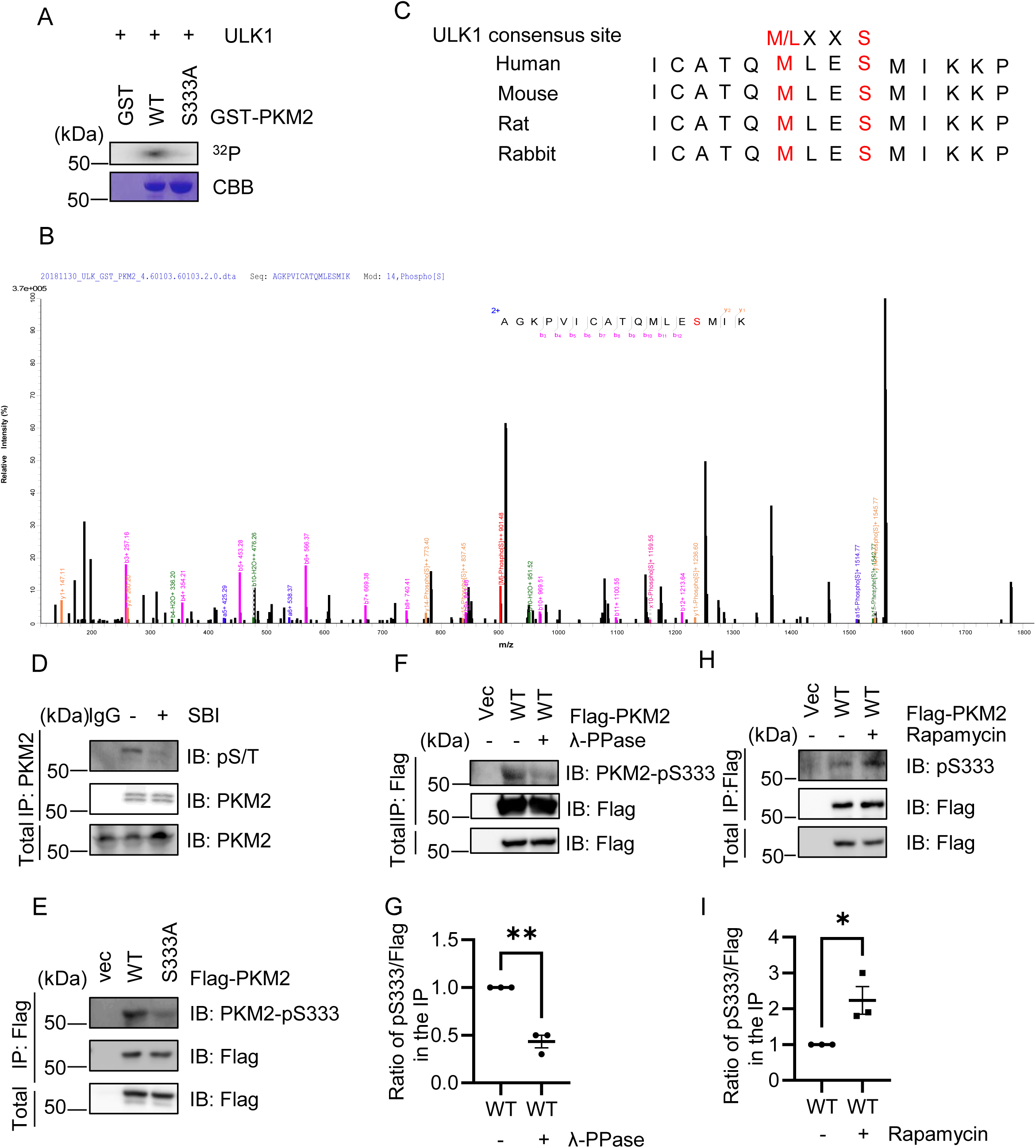
ULK1 phosphorylates PKM2 at pS333. *A,* GST-ULK1 proteins were incubated with recombinant GST-PKM2 or GST-PKM2-S333A in kinase buffers containing ^32^P-ATP. Coomassie Blue (CBB) staining showed the amount of the input PKM2 proteins, and autoradiography detected phosphorylated GST-PKM2. *B*, Mass spectrometry analysis identified a phosphorylated PKM2 peptide. *C*, A comparison of Ser-333 and the ULK1 phosphorylation consensus motif. The residues in red fit the consensus motif. *D*, 293T cells were treated with SBI-0206965 (ULK1 inhibitor) or mock treated; the extracts were then blotted with the antibodies indicated. *E*, Phospho-specificity of PKM2 anti-pS333 antibodies. Cells were transfected with Flag-PKM2-WT and -S333A plasmids, and the extracts were subject to immunoprecipitation with anti-Flag antibodies. *F*, Phospho-specificity of PKM2 anti-pS333 antibody demonstrated by phosphatase (PPase) treatment. Cells were transfected with Flag-PKM2-WT plasmids, and the anti-Flag immunoprecipitates were incubated with λ PPase or left untreated. *G*, Quantitation of F. *H*, Cells were transfected with Flag-PKM2-WT plasmids, then treated with Rapamycin (50 nM, 12 h) or mock treated. *I*, Quantitation of H. The statistical analysis in G and I was performed using Student’s *t*-test, *P < 0.05, **P < 0.01. All Western Blots were performed for at least three times.

To validate that pS333 occurs in cells, we used several methods. First, we used the pan phosphorylation pS/T antibody and SBI-0206965 (SBI) (ULK1 inhibitor). When PKM2 was IPed from cellular lysates (Fig. 2D), SBI treatment markedly decreased the pS/T signals, suggesting that ULK1 mediates PKM2 phosphorylation. Then we generated phospho-specific antibodies targeting PKM2-pS333, whose signal was down-regulated in PKM2-S333A immunoprecipitates and in phosphatase (PPase)-treated PKM2-wild type (WT) immunoprecipitates (Fig.2E-G). Third, Rapamycin was utilized to upregulate autophagy, which significantly increased pS333 levels (Fig. 2H-I). In sum, we found that ULK1-dependent phosphorylation of PKM2 occurs at S333.

### PKM2 S333 phosphorylation antagonizes O-GlcNAcylation by decreasing interaction with OGT

We then set out to validate our hypothesis, whether the glucose sensitive S333 phosphorylation has crosstalk with O-GlcNAcylation. By treating the cells with Thiamet-G (TMG) plus glucose as previously described (20), we enriched the cellular lysates for the O-GlcNAc modification. As shown in Fig. 3A-B, the phospho-deficient S333A mutant increased O-GlcNAcylation, while the phospho-mimic S333D mutant decreased O-GlcNAcylation. We wondered whether the changes in O-GlcNAcylation levels were due to alterations between the binding between PKM2 and OGT, so we tested the interaction between OGT and PKM2-WT and mutants. Exogenous Flag-PKM2-S333A showed a modest increase in binding HA-OGT (Fig. 3C-D). GST-OGT also increased interaction with Flag-PKM2 (Fig.3E-F). When endogenous OGT was assessed, Flag-PKM2-S333A demonstrated a robust increase with OGT (Fig. 3G).

**Fig 3.**
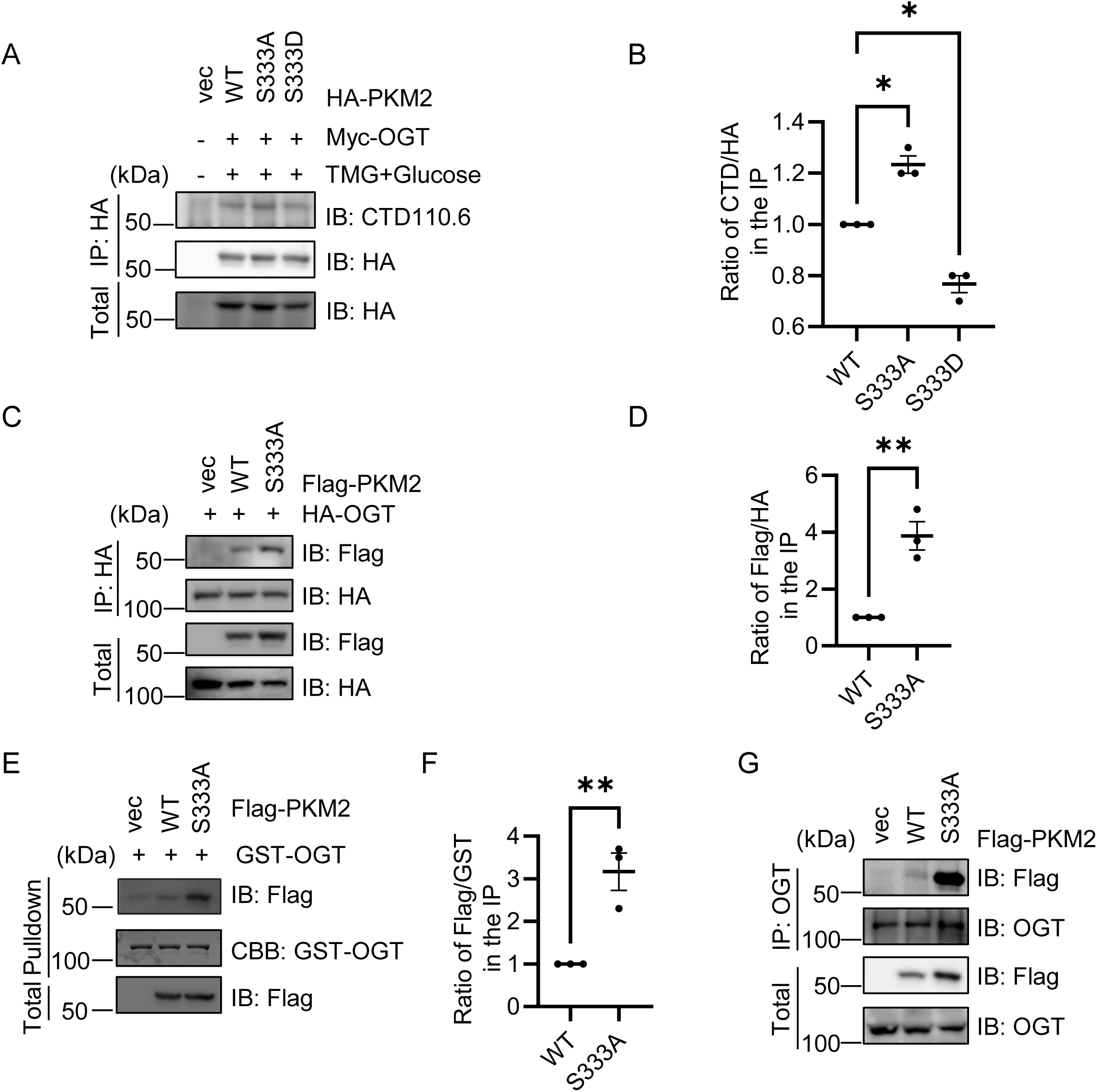
PKM2 pS333 antagonizes O-GlcNAcylation. *A,* 293T cells were transfected with HA-PKM2-WT, -S333A or -S333D plasmids and together with Myc-OGT. The cells were treated with glucose and Thiamet-G (TMG) or mock treated. *B*, Quantitation of A. *C,* Cells were transfected with Flag-PKM2-WT and -S333A and HA-OGT plasmids, and the extracts were subject to immunoprecipitation with anti-HA antibodies and immunoblotted with the indicated antibodies. *D*, Quantitation of C. *E*, Cells were transfected with Flag-PKM2-WT and -S333A plasmids, the extracts were then incubated with recombinant GST-OGT proteins and subject to pulldown assays. *F*, Quantitation of E. *G*, Cells were transfected with Flag-PKM2-WT and -S333A plasmids, and the extracts were subject to IP with anti-OGT antibodies. The statistical analysis in B was performed using one-way Anova, and in D and G using Student’s *t*-test, *P < 0.05, **P < 0.01. All Western Blots were performed for at least three times.

We also examined whether O-GlcNAcylation has crosstalk with pS333 by constructing PKM2-S362A/T365A/T405A/S406A mutants. But the mutant does not show changes in pS333 levels (data not shown). It is possible that PKM2 has other O-GlcNAcylation sites. Molecular dynamics simulations did not reveal any interaction between pS333 and O-GlcNAcylation sites either. These results suggest that PKM2 pS333 antagonizes O-GlcNAcylation, probably due to decreased affinity with OGT.

### PKM2 pS333 decreases pS37 levels and nuclear localization

As PKM2 O-GlcNAcylation has been shown to increase ERK-dependent phosphorylation at S37 (15), we then evaluated the effect of pS333 on pS37. Cells were transfected with PKM2-WT and -S333A plasmids. And pS37 levels were upregulated in the S333A mutant (Fig. 4A-B). The interaction between PKM2 and ERK was then tested, and Fig. 4C showed that PKM2-S333A significantly increased binding with ERK. ERK-mediated phosphorylation of PKM2 is known to mediate its nuclear localization via importin α5, and our biochemical experiments further demonstrated that S333A increased binding with importin α5, which correlates with the results using the ULK1 inhibitor SBI (Fig. 4D-E). Moreover, immunofluorescence microscopy and fractional analysis was carried out to examine PKM2 localization. Both the S333A mutant and SBI treatment increased the nuclear localization of PKM2 (Fig. 4F-I). These results indicate that PKM2 pS333 decreases pS37 by attenuating interaction with ERK.

**Fig 4.**
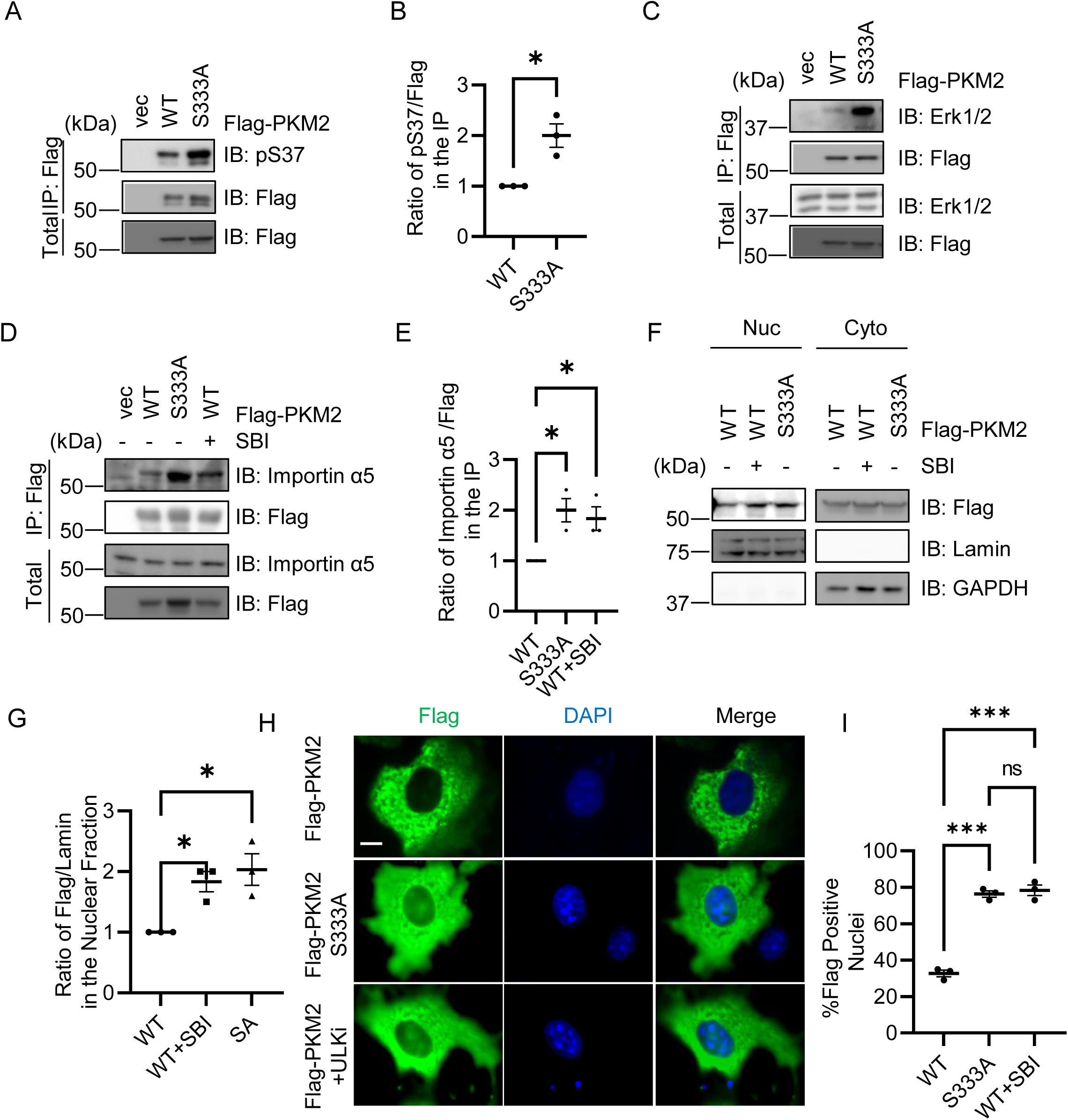
PKM2 pS333 reduces pS37 levels and decreases nuclear localization. *A-C,* 293T cells were transfected with Flag-PKM2-WT and the extracts were then immunoprecipitated with anti-Flag antibodies. The immunoprecipitates were immunoblotted with anti-pS37 antibodies (A) or anti-Erk1/2 antibodies (C). *B*, Quantitation of A. *D,* HEK-293T cells were transfected with Flag-PKM2-WT and -S333A plasmids and treated with SBI-0206965. The anti-Flag immunoprecipitants were immunoblotted with anti-importin α5 antibodies. *E*, Quantitation of D. *F-I*, Cells were transfected with Flag-PKM2-WT or -S333A plasmids and treated with SBI-0206965 or mock treated. Then the cells were analyzed by nuclear cytoplasmic fractionation assays (F-G), or immuno-stained with anti-Flag antibodies and DAPI (H). Immunofluorescence experiments were repeated three times, with 100 cells per experiment. Scale bar, 10 µM. *G,* quantitation of F. *I,* quantitation of H. The statistical analysis in B was performed using Student’s *t*-test, in E, G and I using one-way Anova. *P < 0.05. ***P < 0.001. All Western Blots were performed for at least three times.

### PKM2 pS333 attenuates the Warburg effect

PKM2 O-GlcNAcylation has been shown to promote the Warburg effect, partly through its nuclear role via c-Myc (15). We then tested the effect of pS333 on c-Myc. Stable cell lines were constructed (Fig. 5A). And c-Myc mRNA levels were upregulated in the S333A mutant, as expected (Fig. 5B). The protein levels of c-Myc were also increased (Fig. 5C-D). As the nuclear localization of PKM2 induces c-Myc-mediated glycolysis gene expression such as Glut1 and LDHA (9), we then tested glucose consumption and lactate production (Fig. 5E-F). Both S333A mutants and SBI treatment promoted the Warburg effect.

**Fig 5.**
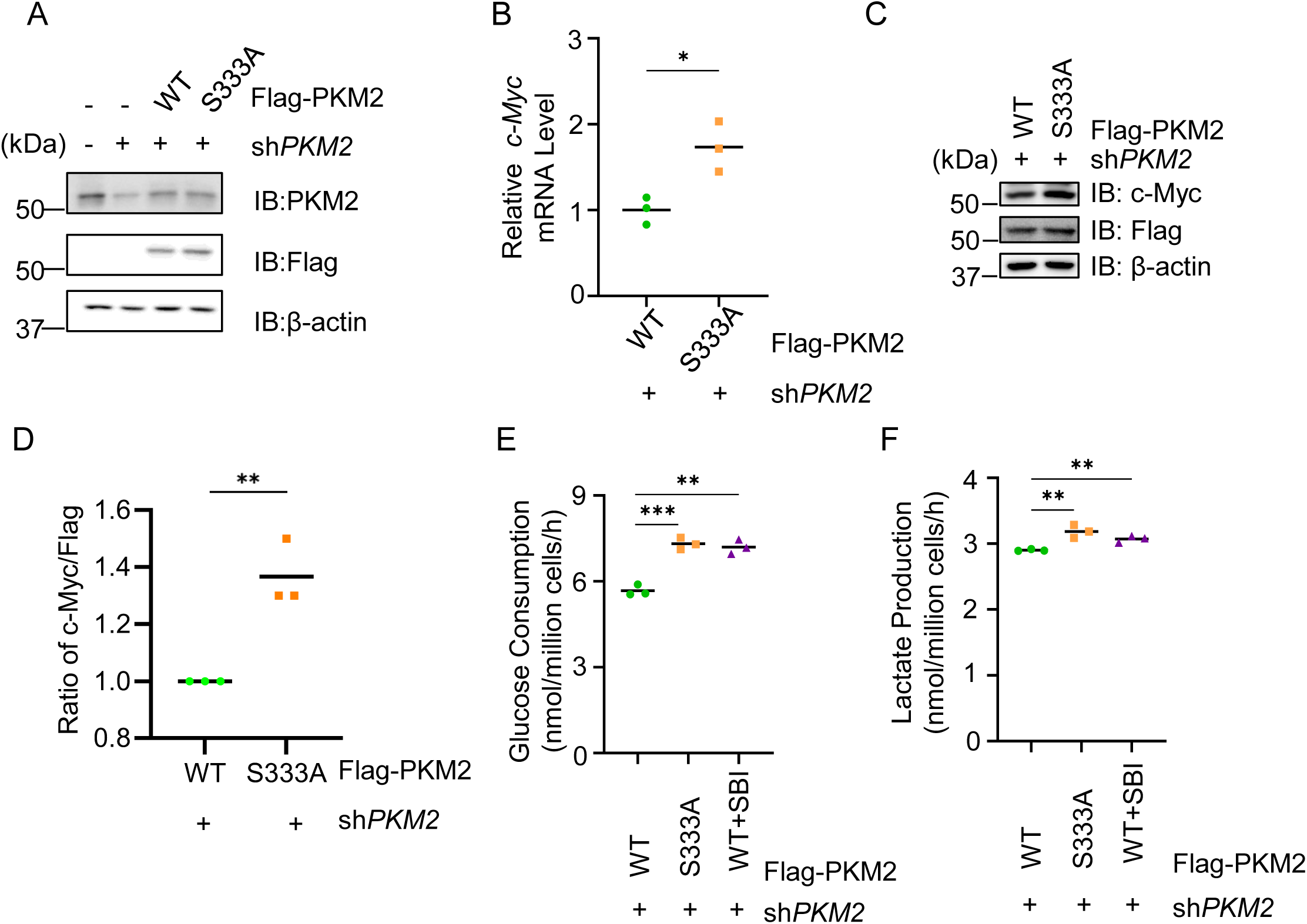
PKM2 pS333 attenuates the Warburg effect. *A,* sh*PKM2* 4T1 cells were transfected with Flag-PKM2-WT and -S333A plasmids, the extracts were then blotted with the antibodies indicated. *B-D,* The cells in A were subject to qPCR to evaluate *c-Myc* expression (B) and c-Myc protein levels (C-D). *D,* quantitation of C. *E-F,* Glucose consumption and lactate production assay. The cells in A were treated with SBI-0206965 or mock treated, then analyzed as described in Experimental procedures. The statistical analysis in B and D was performed using Student’s *t*-test, in E and F using one-way Anova. *P < 0.05, **P < 0.01, ***P < 0.001. All Western Blots were performed for at least three times.

### pS333 promotes PKM2 tetramer formation and enzymatic activity

It is known that PKM2 O-GlcNAcylation destabilizes PKM2 tetramer formation and reduces PK enzymatic activity (15). We then studied the effect of pS333. Fig. 6A showed that both S333A mutants and SBI treatment decreased the tetramer formation of PKM2. As PKM2 tetramers are crucial for its enzymatic activity, PK activity was also evaluated (Fig. 6B), and both S333A mutants and SBI treatment lowered PKM activity. In sum, pS333 promotes PKM2 tetramers and PK activity.

**Fig 6.**
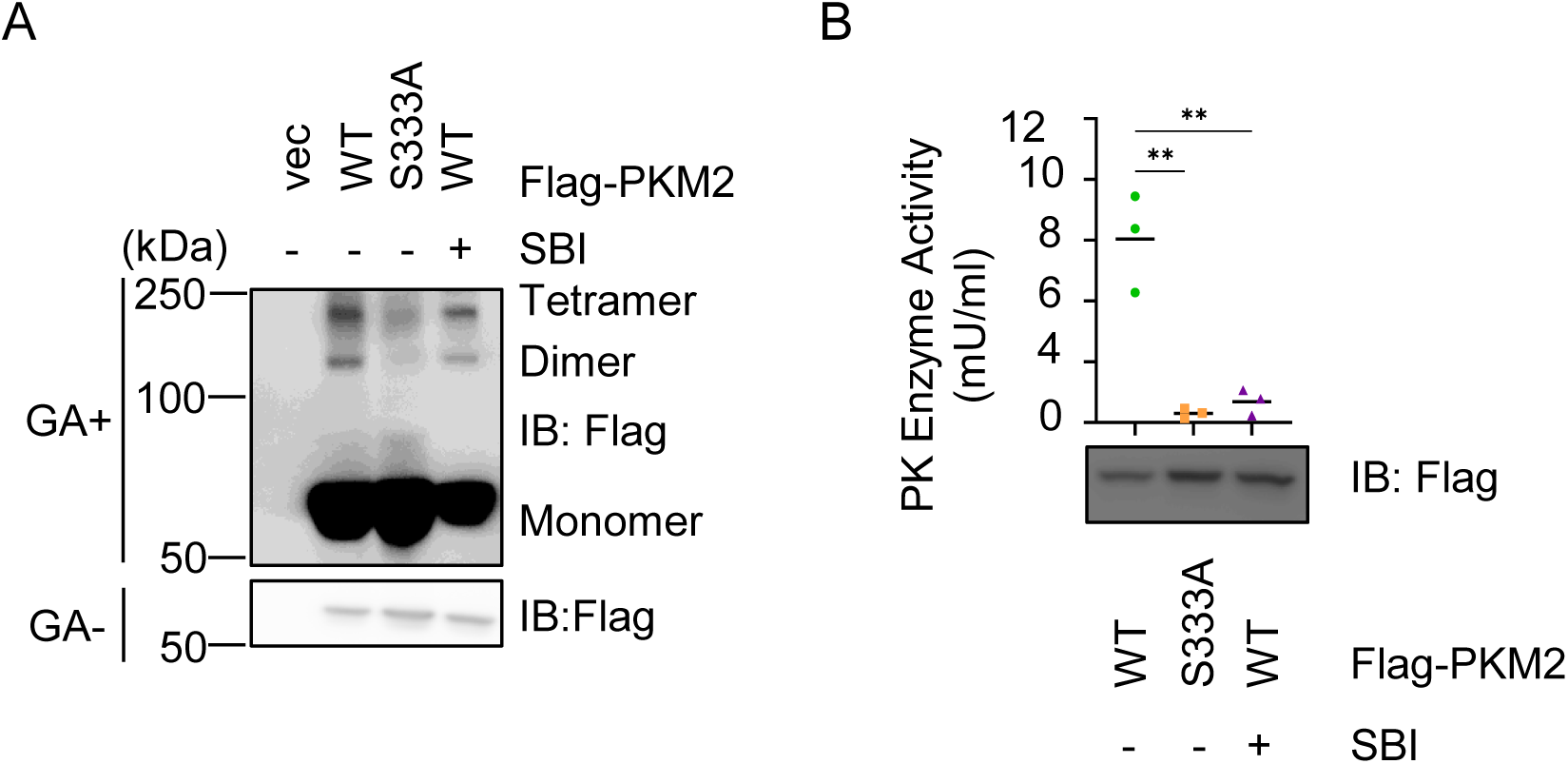
PKM2 pS333 displays more tetramer formation and increases enzymatic activity. *A*, HEK-293T cells were transfected with Flag-PKM2-WT and -S333A plasmids with or without SBI-0206965 treatment. Then PKM2 oligomer states were analyzed as described in Experimental procedures. Whole-cell lysates were analyzed by WB. GA, glutaraldehyde. *B,* Enzymatic activity of PKM2-WT and -S333A was assayed. The statistical analysis was performed using one-way Anova, *P < 0.05, **P < 0.01. All Western Blots were performed for at least three times.

### PKM2 pS333 inhibits breast cancer

To assess the function of PKM2 O-GlcNAcylation *in vivo*, we stably expressed FLAG-PKM2-WT or FLAG-PKM2-S333A plasmids in 4T1 mouse mammary tumor cells (Fig.7A), then injected them subcutaneously into BALB/c mice and observed the tumor growth. PKM2 overexpression promotes tumor growth, while S333A stimulated it further (Fig.7B-7E). The immunohistochemistry analysis revealed that Ki-67-positive tumor cells were significantly increased in both WT and S333A overexpression tumors, while S333A increased it further (Fig. 7F-G). Together, these results support the notion that PKM2 pS333 suppresses tumor growth in xenograft tumorigenesis.

**Fig. 7.**
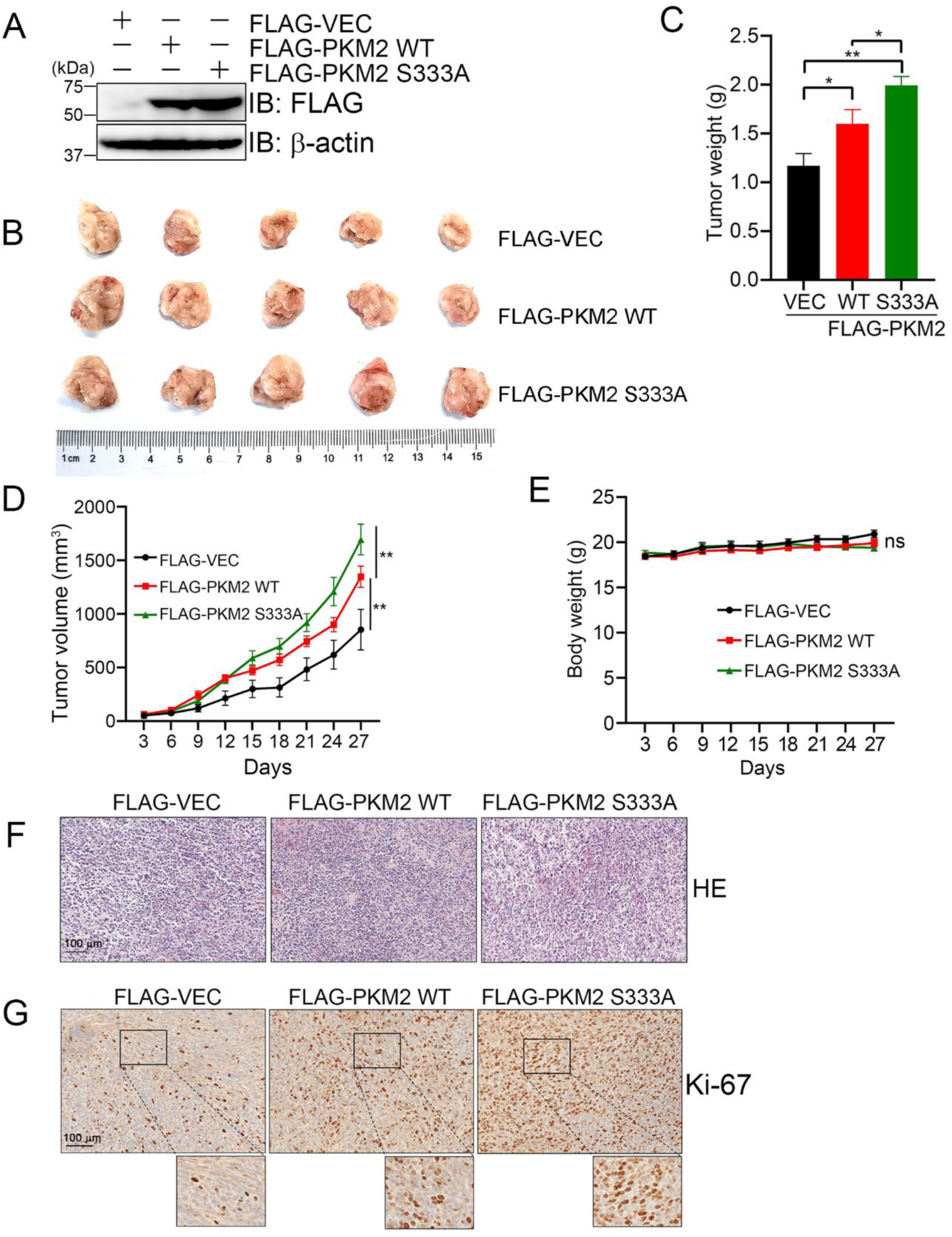
PKM2 pS333 suppresses breast cancer progression. *A*, Immunoblotting analysis of FLAG-VEC, FLAG-PKM2 WT and FLAG-PKM2 S333A in 4T1 murine mammary tumor cells. *B,* The cells in (A) were injected into Balb/c mice (n=5). 27 days post transplantation, the tumors were imaged. *C,* The tumor weight of each group from (B). *D,* Relative volume change of tumors in different groups. *E,* Body weight change in different group. F-G, Histopathological examination of tumors. Representative images of HE staining in tumor samples (F) and representative image of IHC staining showing Ki-67 in tumor samples (G). The statistical analysis in C was performed using one-way Anova, in D and E using two-way Anova, *P < 0.05, **P < 0.01. All Western Blots were performed for at least three times.

**Fig. 8.**
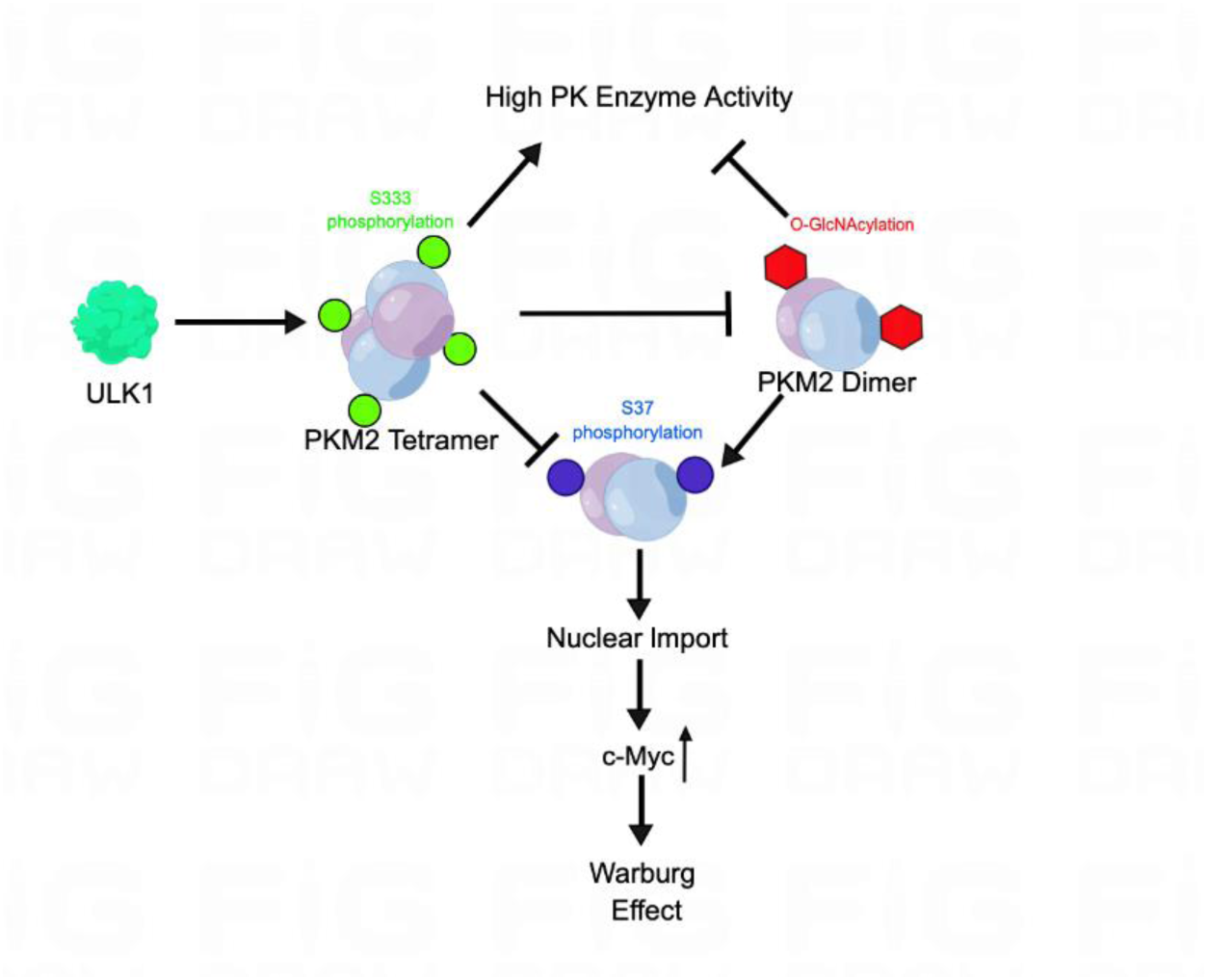
Model depicting the role of PKM2 S333 phosphorylation. ULK1 phosphorylates PKM2 at S333, which antagonizes PKM2 O-GlcNAcylation. PKM2 pS333 inhibits Warburg effect and tumorigenesis.

## Discussions

In this work we present evidence that ULK1 phosphorylates PKM2 at S333, which antagonizes O-GlcNAcylation. pS333 promotes PKM2 tetramer formation and enzymatic activity, and decreases its nuclear localization. PKM2 pS333 attenuates the Warburg effect through downregulating glucose consumption and lactate production, thus inhibiting breast cancer tumorigenesis. Our work identifies a new phosphorylation on PKM2 that is sensitive to glucose concentration.

Previously, PKM2 has been identified to be O-GlcNAcylated at Thr-405 and Ser-406 (15), as well as in Ser-362 and Thr-365 (16). Other PTMs, such as methionine oxidation (12) and ubiquitination (11), also regulate PKM2 activity, localization and oligomerization state. As O-GlcNAc is a nutrient rheostat, it is not surprising that the glucose-sensitive pS333 on PKM2 antagonizes O-GlcNAcylation.

We tried to unearth the mechanism of how pS333 counteracts O-GlcNAcylation by molecular dynamics simulation (data not shown), but the Ser-333 residue is too far away from the O-GlcNAc sites, so any steric effect generated from the modification may not be significant. From our biochemical assay (Fig. 3G), we showed that S333A mutants of PKM2 robustly increased interaction with OGT. But the O-GlcNAc mutants (S362A/ T365A/T405A/ S406A) does not affect pS333 levels or interaction with ULK1, suggesting that pS333 is epistatic to O-GlcNAcylation.

ULK1 is known as the autophagy-initiating kinase, and it redirects energy and resources to autophagy when nutrient is scarce. Our results suggest that as a nutrient-sensing kinase, ULK1 may dictate metabolic reprogramming by mediating PKM2 phosphorylation. We show that pS333 is down regulated when glucose is in abundance, due to reduced interaction between ULK1 and PKM2. As PKM2 is the pivotal glycolysis enzyme, ULK1 thus couples metabolism with the changing environmental stress. As ULK1 is implicated in many human diseases, including neurodegenerative diseases and cardiovascular diseases, our work may provide a new link between the disease pathogenesis and metabolic reprogramming.

In summary, we identified a nutrient-sensitive phosphorylation of PKM2, and showed an interplay between pS333 and O-GlcNAcylation. Various PTMs on PKM2 may crosstalk to finetune its activity to adjust cellular metabolism to the environmental dynamics.

## Experimental procedures

### Cell culture, antibodies and plasmids

HeLa and HEK293 cells were purchased from ATCC. Antibodies were as follows: anti-c-Myc (Abcam, 32072, Cambridge, UK), anti-Erk1/2 (CST, 9102S, Danvers, Massachusetts, USA), anti-Flag(Sigma, F1084, Darmstadt, Germany), anti-GAPDH (TransGen Biotech, H015, Beijing, China), anti-GST (Gene Script, A00865, New Jersey, USA), anti-HA (Bethyl, A190-108A, Texas, USA), anti-IgG Rabbit (Sigma, R2665, Darmstadt, Germany), anti-Importin α5 (SAB, 25191, Greenbelt, Maryland, USA), anti-Lamin A/C (CST, 2032S, Danvers, Massachusetts, USA), anti-O-GlcNAc (CTD110.6), (Sigma, 078M4878V, Darmstadt, Germany), anti-OGT (Santa Cruz Biotechnology, sc-74546, Santa Cruz, CA, USA), anti-pS/T (ECM Biosciences, PP2551, New Jersey, USA), anti-PKM2 (CST, 4053S, Danvers, Massachusetts, USA), anti-PKM2-pS37 (SAB, 11456, Greenbelt, Maryland, USA), anti-ULK1 (CST, 8054, Danvers, Massachusetts, USA) and anti-β-actin (Sigma, A5441, Darmstadt, Germany). Anti-PKM2-pS333 antibodies targeting MLEpSMIK were manufactured by Dia-An Biotech, Inc. (Wuhan, China). shRNA sequences targeting PKM2 were: shPKM2: CAACGCTTGTAGAACTCACTC. Mutations (including PKM2-S333A and PKM2-S333D) were made by using the KOD-Plus-Mutagenesis Kit (TOYOBO, SMK-101, Osaka, Japan).

### IP and immunoblotting

The following primary antibodies were used for IB: anti-c-Myc (1:4000), anti-Erk1/2 (1:1000), anti-Flag (1:4000), anti-GAPDH (1:5000), anti-HA (1:3000), anti-Importin α5 (1:1000), anti-Lamin A/C (1:1000), anti-O-GlcNAc (CTD110.6) (1:1000), anti-OGT (1:1000), anti-Phosphoserine/threonine (1:1000), anti-PKM2 (1:1000), anti-PKM2-pS37 (1:1000), anti-ULK1 (1:1000), anti-β-actin (1:5000), anti-PKM2-pS333 (1:1000). Peroxidase-conjugated secondary antibodies were from JacksonImmuno Research. The ECL detection system (Amersham, Bath, UK) was used for immunoblotting. LAS-4000 was employed to detect signals, and quantitated by the Multi Gauge software (Fujifilm). All western blots were repeated for at least three times.

### Chemical treatment

Thiamet-G (Sigma, Darmstadt, Germany) was used at 0.25 mM for 24 h; glucose was used at 30 μM for 3 h.; SBI-0206965 (MCE, New Jersey, USA) was used at 10 μM for 3 h.

### Immunofluorescence Microscopy

Immunofluorescence staining was carried out as described previously (18). Antibody dilutions were 1:1000 for mouse anti-Flag. The nuclei were stained with DAPI. All immunofluorescence experiments were repeated three times, with 100 cells per experiment.

### PKM2 oligomerization assay

The whole-cell lysates (4 mg/mL) were cross-linked with 0.025% glutaraldehyde for 3 min at 37 °C and terminated with Tris-HCl (pH 8.0, 50 mM). Subsequently, samples were analyzed by Western blotting with indicated antibodies.

### RNA extraction and quantitative real-time PCR

After the cell lysates were collected, RNAi So Plus (Takara Bio Inc, #9108Q, Osaka, Japan) was added to resuspend the cells. After centrifugation at 12,000 rpm for 15 min, the supernatant was taken and mixed with isopropanol. After centrifugation at 4 °C at 12,000 rpm for 10 min, the supernatant was discarded and 250 μL RNase-Free Water was added to dissolve the precipitate. After adding anhydrous ethanol and mixing, the mixture was centrifuged at 4 °C at 7500 g for 5 min. The supernatant was discarded, and the precipitate was dried at room temperature. RNA was then dissolved using RNase-Free water. The extracted RNA was stored in a refrigerator at -80 °C. After measuring the concentration, the cDNA reverse transcription process was performed. The AceQ-qPCR SYBR Green Master Mix kit (Vazyme, #Q111-02, Nanjing, China) was used for further analysis.

### Glucose consumption and lactate production assay

Media from cultured cells was collected for the assay of glucose and lactate. Glucose levels were determined by using a glucose assay kit (Sigma, # GAGO20-1KT, Darmstadt, Germany). Glucose consumption was the difference in glucose concentration compared with DMEM. The lactate level was determined by using fluorescence-based assay kits (Bio Vision, # K607, Cambridge, UK) according to the manufacturer’s protocols.

### In vitro PK Enzymatic activity assay

Cells were transfected with Flag-PKM2 and -S333A plasmids. Then the Pyruvate Kinase Activity was detected by using the Pyruvate Kinase Activity Colorimetric Assay Kit (Bio Vision, K709, Cambridge, UK).

### In vitro phosphatase assay

HEK 293T cells were transfected with Flag-PKM2 plasmids. The anti-Flag immunoprecipitates were subject to Lambda phosphatase treatment (NEB #P0753L, Ipswich, MA, USA).

### In vitro kinase assay

GST-PKM2 and GST-PKM2-S333A proteins purified from *E. coli*. They were incubated with recombinant ULK1 proteins (Sigma, #SRP5096-10UG, Darmstadt, Germany) in the kinase buffer (20 mM HEPES (pH 7.5), 1 mM EGTA, 0.4 mM EDTA, 5 mM MgCl2, 0.05 mM Dithiothreitol, 0.1 mM ATP or 1 mCi of ^32^P-ATP at 37[for 1 h. The reactions were stopped by boiling at 100[with 6 X sample buffers.

### Mass spectrometry (MS)

For identification of phosphorylation by MS, proteins isolated by gel electrophoresis were digested with trypsin (Promega, Wisconsin, USA) in 100 mM NH_4_HCO3 (pH 8.0). The LC-MS/MS analysis was performed on an Easy-nLC 1000 II HPLC (Thermo Fisher Scientific, Waltham, MA, USA) coupled to a Q-Exactive HF mass spectrometer (Thermo Fisher Scientific, Waltham, MA, USA). Peptides were loaded on a pre-column (100 μm ID, 6 cm long, packed with ODS-AQ 10 μm, 120 Å beads (YMC Co., Ltd. Kyoto, Japan) and further separated on an analytical column (75 μm ID, 15 cm long, packed with Luna C18 1.9 μm 100 Å resin from Welch Materials)(West Haven, CT, USA) using a linear gradient from 100% buffer A (0.1% formic acid in H_2_O) to 30% buffer B (0.1% formic acid in acetonitrile), 70% buffer A in 80 min at a flow rate of 200 nL/min. The top 20 most intense precursor ions from each full scan (resolution 120,000) were isolated for HCD MS2 (resolution 15,000; normalized collision energy 27) with a dynamic exclusion time of 60 s. Precursors with a charge state of 1+, 7+ or above, or unassigned, were excluded.

The software pFind3 (21) was used to identify phosphorylated peptides by setting a variable modification of 79.966331 Da at S, T, and Y, and a neutral loss of 97.976896 Da at S and T. The mass accuracy of precursor ions and that of fragment ions were both set at 20 ppm. The results were filtered by applying a 1% false discovery rate cutoff at the peptide level and a minimum of one spectrum per peptide. The MS2 spectra were annotated using pLabel (22).

### Mouse Xenograft Analysis

Female BALB/c mice, aged 4 to 6 weeks, were purchased from Weitong Lihua Laboratory Animal Technology Co., Ltd. (Beijing, China). The mice were randomly divided into three groups. The xenograft model was established by injecting about 5*10^5^ 4T1 murine mammary tumor cells which stably expressing FLAG-VEC, FLAG-PKM2 WT and FLAG-PKM2 S333A subcutaneously into the right leg of mice (n=5 in each group). Tumor size was measured using Vernier calipers. When the tumor volume reached 1500 mm^3^, the mice were killed and the tumor were removed followed by HE and IHC. The guidelines for the care and use of laboratory animals were followed for all animal experiments, and approved by the animal care committee of Shenzhen University.

### HE staining and immunohistochemistry (IHC) assay

The isolated mice tumor tissues were fixed in 4% paraformaldehyde, embedded in paraffin and cut into 2-mm sections. The sections were subsequently deparaffinized in 60 □ for 2 hours and stained with hematoxylin and eosin (HE). For IHC assay, the deparaffinized sections were blocked with 5% BSA at 37 □ for 30 min and incubated with antibody against Ki-67 (Ki-67 (8D5) Mouse mAb #9449) overnight at 4 □. Subsequently, immunostaining was performed according to the standard protocol of the DAB Substrate Kit (ZSGB-BIO, PV-6000). Images were captured with a Pathology Slice Scanner (LEICA-Aperio CS2) at ×10 and ×20 magnification.

## Data Availability Statement

The mass spectrometry proteomics data have been deposited to the ProteomeXchange Consortium (http://proteomecentral.proteomexchange.org) via the iProX partner repository (23) (24) with the dataset identifier PXD044583.

## Abbreviations

(MS): Mass spectrometry
(O-GlcNAc): O-linked β-N-acetylglucosamine
(OGT): O-GlcNAc transferase
(PTM): post-translational modification
(IP): Immunoprecipitation
(IB): Immunoblotting
(PKM2): Pyruvate kinase M2
(ULK1): unc-51 like autophagy activating kinase 1
(SBI): SBI-0206965

## Acknowledgements

We thank Dr. Hong-Jie Zhang (Core Facility for Protein Research, Institute of Biophysics, Chinese Academy of Sciences) for the technique support using radioactivity detection. J. L. is supported by the National Natural Science Foundation of China (NSFC) fund (32271285 and 31872720) and R & D Program of Beijing Municipal Education Commission (KZ202210028043). X. X. is supported by NSFC fund 32090031, the Shenzhen Science and Technology Innovation Commission projects JCYJ201805073000163.

